# ProteoClade: a taxonomic toolkit for multi-species and metaproteomic analysis

**DOI:** 10.1101/793455

**Authors:** Arshag D. Mooradian, Sjoerd van der Post, Kristen M. Naegle, Jason M. Held

## Abstract

We present ProteoClade, a Python toolkit that performs taxa-specific peptide assignment, protein inference, and quantitation for multi-species proteomics experiments. ProteoClade scales to hundreds of millions of protein sequences, requires minimal computational resources, and is open source, multi-platform, and accessible to non-programmers. We demonstrate its utility for processing quantitative proteomic data derived from patient-derived xenografts and its speed and scalability enable a novel *de novo* proteomic workflow for complex microbiota samples.

## Main Text

The goal of metaproteomic and multispecies proteomic studies is to characterize the proteomes of samples containing multiple, comingled species, which can provide insight into the complex interactions at the interface between organisms. Proteomic analysis of these samples can quantify thousands of proteins from hundreds of species in a single mass spectrometry experiment^1^, characterize education of stromal tissue by patient-derived xenografts (PDXs)^2^, and extensively characterize the human oral microbiome^3^.

Metaproteomic and multispecies data analyses depend on the ability to integrate reference protein sequence databases, taxonomic lineages, *in silico* proteolytic digestion, peptide identification, and quantitation. These studies universally perform bottom-up analysis, where proteins are digested into peptides with a protease, and therefore require assignment of peptides to proteins based on their taxonomic specificity. Several software tools provide one or more of these features, but have practical and technical limitations that render them unable to facilitate complete analysis pipelines of quantitative proteomics data and scale to the rapidly increasing number of available reference protein sequences^4,5^. With regard to annotating peptides to taxa, Unipept is a commonly used taxonomic annotation tool that can provide access to the entire UniProt sequence repository, provides web-based visualizations and a command line interface, and was demonstrated to annotate peptides orders of magnitude faster than a prior UniProt-based application, Peptide Match^6,7^. However, the Unipept database is unavailable for the end-user to customize, is restricted to a fixed set of assumed experimental parameters such as protease, cannot be used with custom protein databases such as those generated by sequencing, and lacks capability for protein quantitation which limits its utility for analyzing many experimental data sets. Additionally, the Unipept database was generated using high performance computing resources which poses a technical challenge as the number of sequences in UniProt grows exponentially^5,8^. Other tools offering more complete metaproteomic pipelines such as MetaProteomeAnalyzer (MPA) support database generation, peptide spectral matching, and taxonomic parsing. However, these tools restrict the user to bundled open-source targeted database search engines which scale poorly when using large reference database sizes such as the entirety of UniProt^9^. MPA also lacks support for post-translational modifications and common MS2-based quantitation approaches, limiting its applicability.

To overcome these limitations, we developed ProteoClade, an open-source Python library that enables flexible, rapid, and easy taxon-specific quantification for proteomic experiments. ProteoClade utilizes standard taxonomic and protein sequence repositories and is optimized for large databases. ProteoClade is the first tool to 1) enable users to generate and search customizable, *in silico* digested peptide-to-taxa mapped databases that can scale to the entire UniProt database with the optional inclusion of user-specified reference protein sequences; and 2) provide a novel *de novo* workflow to efficiently annotate peptides sequenced without defining the taxonomic composition *a priori*. Additionally, ProteoClade allows the user to choose their preferred commercial or open-source search engine, as well as preserves MS1-, MS2-, and spectral count-based quantitation from the experiment to calculate gene-level, taxon-specific quantitative results. Together, ProteoClade uniquely enables fast, taxon-specific quantitation of database-targeted multispecies experiments as well *de novo* searches enhanced by large databases for complex metaproteomic experiments. The ProteoClade software is freely available at http://github.com/HeldLab/ProteoClade.

ProteoClade retrieves complete taxonomic lineages from the NCBI, and interfaces directly with the UniProt API to download and concatenate reference protein sequence databases based on the organism IDs and database parameters supplied by the user. To assign peptide-spectral matches from mass spectra search engines to both taxon and gene identifiers, ProteoClade creates a SQLite database (ProteoClade Database: PCDB) by digesting reference proteomes *in silico* with proteolytic parameters that can be customized according to the user’s experimental conditions (**Fig. S1**). The PCDB addresses scalability issues by storing peptide sequences as hashed integer values which compresses the average peptide storage requirement by 62.8%. ProteoClade leverages multiple CPUs for parallel processing to speed both the PCDB creation and search functions, and indexes the database to quickly assign peptides to taxon-specific genes. ProteoClade’s implementation is further detailed in the STAR methods.

Metaproteomic experiments involving communities of many organisms present substantial computational challenges since more sequence information available to aid in the identification and quantitation of the experimental data requires more computational resources. Thus, it is imperative to efficiently store and parse these data for taxonomic assignment and protein quantitation in light of the dramatic increase in the number of organisms with proteomic annotation. We evaluated ProteoClade’s scalability by creating a tryptic PCDB from the November 2018 release of UniProt containing 140.2 million protein entries (UniProt PCDB). The resultant database contained 10.7 billion peptides, 5.02 billion of which were unique, from 1,040,460 organisms. PCDB creation, including indexing, took only 11 hours on modest consumer-grade hardware and resulted in a 515GB file (**Fig. S2a**), demonstrating that ProteoClade enables the use of large, customizable peptide databases.

We compared the database indexing and taxonomic annotation features of Unipept, MetaProteomeAnalyzer (MPA), and ProteoClade to highlight ProteoClade’s optimizations and improvements over prior taxonomic tools. For peptide database creation and indexing, ProteoClade uses 32-fold less RAM and 6-fold less time than Unipept, and > 60-fold less RAM and > 120 fold less time than MPA (**Fig. S2a, S2b**). We found that ProteoClade annotates experiments at 8.8x the speed of Unipept and preserves quantitative information with the ability to sum the peptides’ quantitation to the gene level as is common in multispecies experiments, while MPA lacked the ability to annotate peptides outside of database-targeted searches (**Fig. S2c**). ProteoClade’s technical optimizations enable taxonomic annotation and quantitation of peptides at a speed and scale that exceed previously used tools.

To demonstrate ProteoClade’s ability to perform integrated, species-specific quantitation of multi-species samples, we analyzed publicly available TMT-labeled global proteomics data from Patient-derived xenograft (PDX) lines in which six triple-negative breast cancer tumors were each grafted into three mice, but were originally analyzed without considering mouse-specific peptides^10^. PDXs are a burgeoning, mixed-species model of tumor biology^11^ in which the stromal microenvironment of tumors, comprised of fibroblasts, immune-related cells, and vasculature, is originally human but is replaced after several passages by murine cells. Thus, species-specific proteomic analysis of PDX data can simultaneously and independently characterize the tumor and the invasive murine stroma to examine how tumors remodel stromal proteomes in a process known as stromal education^2^.

ProteoClade was used to create a customized, concatenated human and mouse UniProtKB/Swiss-Prot database for both peptide-spectral matching with MaxQuant and creation of a PCDB for species-specific taxonomic annotation and filtering (**Fig. 1a**). In addition, ProteoClade’s quantitation module flexibly allows for species-specific summation of TMT reporter ions to each peptide’s assigned gene symbol to generate quantitative proteomic maps of the tumor and stromal proteomes.

**Figure 1:**
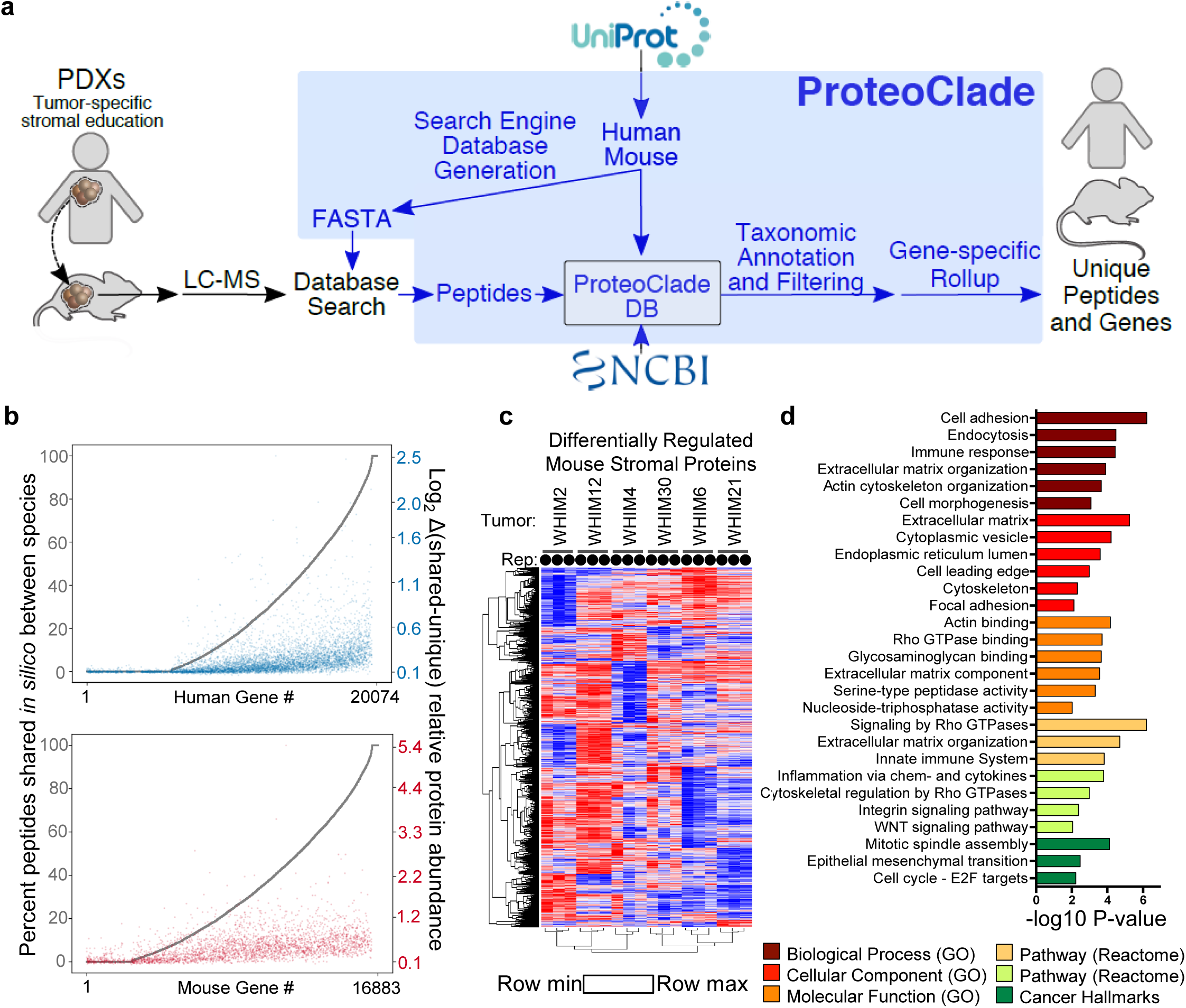
ProteoClade easily interrogates the interaction between comingled species, including PDX tumors’ education of their microenvironment. **a)** The ProteoClade workflow for targeted database searches. ProteoClade downloads and concatenates reference FASTA sequence databases and builds a ProteoClade database for fast taxon-specific protein quantitation. **b)** Comparison between taxon-specific and taxon-shared data processing approaches for a PDXid data setid. The theoretical fraction of shared peptides for each proteome (grey) correlates with the observed quantitative difference between the approaches. A baseline difference between these two analytic approaches was present due to how the assumptions made when assigning peptides to genes affected data normalization. **c)** A significant portion of the murine stromal proteome is differentially regulated by tumors. 3 biological replicates ‘Rep’ per tumor, indicated with circles. **d)** Pathway enrichment analysis of differentially regulated stromal proteins indicates an enrichment in proteins involved in cellular adhesion and the immune response.

Taxon-specific peptide assignment with a tool such as ProteoClade is important in multi-species samples since peptides with shared amino acid sequences between human and mouse may bias proteomic quantitation of PDXs. By digesting peptides *in silico*, we found that 71.1% of genes in the human proteome produce tryptic peptides with sequences identical to their murine homolog, and thus most human and mouse proteins are potentially susceptible to quantitative interference by the presence of the other organism. We examined the consequences of normalizing and quantifying the human and mouse components of the data by comparing a naive informatic assumption that only one organism was present to utilizing ProteoClade for species-specific peptide filtering and protein quantification (**Fig. 1b**). The relative protein abundance varied by more than 1.5-fold for 262 genes in the human-specific data and 891 genes in the murine-specific data, highlighting the large bias that the inclusion or exclusion of species-shared peptides can have on the quantitation. Overall, genes share a higher proportion of tryptic peptides have larger differences in gene-level quantitation indicating that taxonomic interference has a substantial effect on the resulting data (**Fig. 1b**).

ProteoClade greatly facilitated species-specific quantitation of this PDX dataset^10^, identifying 2,326 murine proteins in the microenvironment that are significantly altered by patient-derived tumors, the most expansive set of tumor-educated stromal proteins to date. The murine proteins clustered by biological replicate, confirming that tumor-intrinsic factors drive tumors to persistently educate the stromal proteome as previously observed (**Fig. 1c**)^2^. ANOVA revealed that stromal education by tumors is wide-spread, with 77.1**%** of stromal proteins differentially regulated by the embedded tumors. Differentially-regulated stromal proteins combined from this analysis and a prior study (n = 3,015 proteins) were enriched for proteins in the extracellular matrix, cytoskeletal processes, and myeloid-derived immune components which are important contributors to tumor growth and metastasis (**Fig. 1d**)^2,10^.

Beyond two-species systems, we examined ProteoClade’s applicability to large-scale metaproteomics workflows using the entire UniProt repository, which enables taxonomic annotation to all potential organisms in the sample (**Fig. 2a**). One important consideration for metaproteomic analysis is that all sequence databases are biased by the inclusion and exclusion criteria selected by their curators. Overrepresentation of peptidomes from certain genera and peptide sequence similarity between related organisms can result in misattributing taxon-specific peptides due to errors in mass spectrometry-based detection and sequencing. We establish that UniProt has a substantial sequence bias from the perspective of bottom-up proteomics experiments by quantifying the number of peptides contained in the UniProt PCDB for each genus as well as the proportion of peptide sequences that were unique (**Fig. 2b**). By mapping all 5.02 billion unique peptides back to their respective genera, we found that taxonomic redundancy from the inclusion of thousands of bacterial strains results in vast overrepresentation of tryptic peptides from several genera, including *Streptomyces, Pseudomonas*, and *Bacillus*. These overrepresented genera are likely to be identified in nearly every genus-specific proteomic analysis regardless of their presence or absence in the sample, highlighting the importance of user-customization to database generation. We provide a table of these data to help inform when selecting a narrower database scope for certain genera in UniProt may be appropriate (**Table S1**).

**Figure 2:**
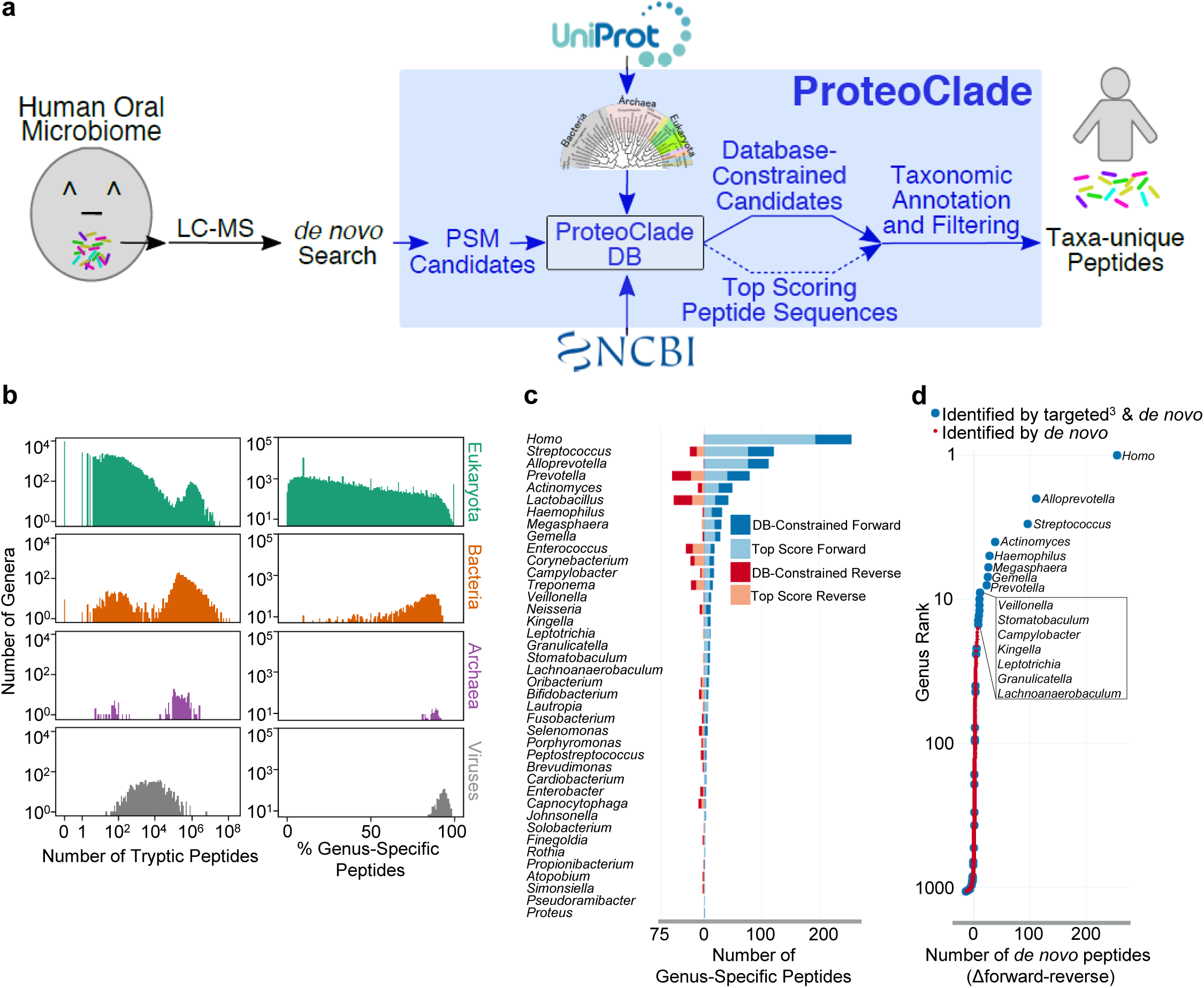
ProteoClade enables identification of taxa-specific peptides in metaproteomic samples by *de novo* sequencing. **a)** The schematic for ProteoClade’s *de novo* pipeline supports the entire UniProt database and the conversion of peptide-spectral candidates to taxon-specific peptides. **b)** A characterization of the entire UniProt peptidome reveals the overrepresentation in the number of peptides contributed by bacterial genera (left), and the significant peptide diversity of microorganisms present in the repository (right). **c)** 39 bacterial genera from the Human Oral Microbiome Database were identified by the *de novo* search in addition to human. DB-Constraned: ProteoClade enhances *de novo* searches by providing the ability to constrain sequence candidates to a database for biological plausibility. “Reverse”: ProteoClade controls the false discovery rate by generating a reversed sequence database. **d)** Comparing the forward and reverse UniProt databases resulted in human and 14 oral bacterial genera being identified by the *de novo* approach. Additional organisms were annotated but these annotations lacked statistical confidence.

*De novo* proteomic analysis is advantageous for metaproteomics since it does not require pre-specifying which organisms are present, which is a requirement for the database-targeted MS2 search approaches universally utilized in metaproteomics. ProteoClade integrates with *de novo* searches since it can taxonomically annotate and quantify *de novo* search results in a unique workflow that can identify organisms and proteins without *a priori* specification, obviating the need for 16s rRNA sequencing or metagenomic assembly for proteomics studies. ProteoClade enables this unique informatic workflow by assigning millions of candidate peptide sequences from *de novo* searched spectra unbiasedly to all possible organisms, which ProteoClade makes possible without the use of high performance computing resources (**Fig. S2**).

We evaluated using ProteoClade for *de novo* peptide assignment of a large oral microbiome proteomic data set^3^ previously analyzed with a standard database-targeted approaches by coupling the UniProt PCDB with *de novo* MS2 assignment (**Fig. 2a**)^3^. ProteoClade parsed 8.13 million peptide-spectral match candidates to annotate 1.83 million MS/MS scans in 1.94 hours (261 spectra per second), making this time-efficient even when challenged with large data sets and reference databases. We identified genus-unique peptides for 39 genera from the oral microbiome in this dataset without prior specification of the bacterial taxa present in the sample, despite the fact that our criteria for identification included genus-specificity in the context of more than 72,000 genera compared to the 108 genera present in the Human Oral Microbiome Database used for the original publication (**Fig. 2c**)^12^.

We next developed an approach to serially check every candidate sequence for each MS2 spectrum until a match in the PCDB was found, and compared these results to an approach which only considers the top scoring candidate for each spectrum. As *de novo* search engines lack a reference database and do not provide false discovery correction for peptide sequencing, we additionally used ProteoClade to generate a reverse ‘decoy’ sequence PCDB, and annotated the peptide sequence candidates using the parameters we chose for our forward search. We then compared the proportion of annotations to the forward database to the reverse database for each genus, which is the same strategy that is used for most targeted database searches, but is novel to *de novo* searches^13^. The combination of considering multiple candidate amino acid sequences for every spectrum and offsetting false discovery with a decoy database increased the number of identified genus-unique human peptides by 32.6% in the forward PCDB annotations, while we observed no increase in the reverse, decoy PCDB annotations (**Fig. 2c)**. This demonstrates that ProteoClade can identify more biologically-valid peptide sequences than would otherwise be offered by upstream *de novo* search software. For several bacterial genera, including *Prevotella* and *Lactobacillus*, we found an increase in both the forward and reverse PCDB annotations, indicating that while the number of peptide candidate sequences increased, those candidates are likely false positives. By comparing all forward and reverse annotations, we observed a threshold (>7 Δforward-reverse genus specific peptides) above which only genera known to have been identified in the human oral cavity are present. We identified human and 14 bacterial genera above this threshold without prior specification, including *Streptococcus*, which was previously reported as the most abundant genus in the oral microbiome (**Fig. 2d**)^14^.

ProteoClade enables taxon-specific analysis of targeted database and *de novo* proteomic experiments. It functions on all major operating systems and operates at a speed and scale that enable fast and novel forms of proteomic data processing using consumer hardware. We expect ProteoClade’s ease of use, applicability to a broad set of biological model systems such as PDXs and metaproteomics, and its novel integration with *de novo* spectral searches^4^ provides a unique and powerful tool to researchers performing quantitative proteomic analysis of multiple mixed species.

## Materials and Methods

### ProteoClade implementation and testing

ProteoClade was developed using Python 3.6 (64-bit) and has been tested on Windows 7, MacOS High Sierra (10.13), and Ubuntu Linux (18.04). All functions have been run and timed using a computer with an Intel i7-2600k processor, 16 gigabytes of RAM, and a Samsung 860 EVO solid state drive. Taxonomic rankings are retrieved directly from the NCBI FTP servers and are assembled as a pickled Python dictionary object which is stored in RAM and allows the software to rapidly assign higher level taxonomies from organism taxonomy cross-reference (OX) IDs with O(1) time complexity. Protein sequences are retrieved using UniProt’s REST API to enable the selection of specific combinations of taxa or database sources (SwissProt, TrEMBL, and/or Reference) using the OX IDs.

The ProteoClade Database (PCDB) and all analyses we performed used the default digest parameters of peptides that range from 7-55 amino acids in length, trypsin with C-terminal proline allowed (trypsin/p), protein N-terminal methionine excision, leucine-isoleucine interconversion, and all combinations up to two missed cleavages; however, we enable the user to select from any variation of these rules, including alternative built-in or custom proteases and alternative ranges of amino acid lengths. Peptides are stored as hashed integers in the PCDB in order to reduce the on-disk storage size. The PCDB creation process was optimized with the use of multiprocessing for computing the hashed peptide integers and integrating them into the database. Similarly, peptide indexing makes use of multiple threads which reduces the total time required for database creation, and is saved as a separate step at the end of database creation to allow the database creation time to scale linearly with the number of protein sequences inserted into it.

Peptide annotation was implemented as a multithreaded process and pushes the bottleneck of ProteoClade’s performance to the input-output operations per second (IOPS) of the harddrive. Annotation can be performed at any combination of taxa the user requests. Quantitative information from search engine outputs is preserved during annotation, and is assigned to gene symbols on the basis of filters supplied by the user. These filters include specifying a taxonomic level to determine the taxonomic uniqueness of a sequence, inclusion and exclusion lists for taxa that the use may want to include or exclude, and a default taxon that can be used to assign quantitative information in the event that multiple taxa are present but only one is of interest to the user, all of which allow the user to create taxon-specific data sets with flexible criteria. Detailed documentation for user-facing functions is included at https://proteoclade.readthedocs.io.

ProteoClade was benchmarked against MetaProteomeAnalyzer (MPA Portable v. 1.9) and Unipept (v. 1.4.1) to assess database creation and taxon annotation speeds. Benchmarking hardware was the same as described above for ProteoClade’s testing. For MPA, time and RAM requirements for peptide indexing were monitored over a 24 hour period of attempting to search a simple mass spectra file using the November 2018 UniProt repository, and figures listed in Fig. S2 are based on the projected estimates, as the software had consumed 14 GB out of 16 GB available RAM in the first 24 hours while only processing the first 2 million (out of 140 million) protein sequences. Unipept times and RAM requirements are taken from the latest Unipept publication^1^ as the Unipept team reported using a high performance computer, and additionally database creation is not a feature of the software that is available for end users to perform. For taxa annotation speed, three files of 5,000 peptides were randomly drawn from the database, and were taxonomically annotated using Unipept’s “pept2taxa” and ProteoClade’s “annotate_peptides” functions.

### PDX proteomic search

Raw spectra files for the Mundt et al. study^2^ were downloaded from the Clinical Proteomics Tumor Analysis Consortium (CPTAC) data portal (https://cptac-data-portal.georgetown.edu/cptac/public). ProteoClade was used to retrieve and concatenate the January 2018 human and mouse SwissProt proteome references into a single FASTA file. A PCDB was generated using the same concatenated database with default settings to enable quick annotation. Spectra were searched using MaxQuant 1.6.0.16 with the following parameters: the instrument acquisition settings were set to the default Orbitrap parameters and the protease selected was trypsin/p. Cysteine carbamidomethylation was set as a fixed modification, with protein N-terminal acetylation and methionine oxidation set as variable modifications. TMT 6-plex MS2 ions were used for quantification. The PSM and peptide FDRs were set to 0.01.

### ProteoClade PDX annotation and quantitation

ProteoClade was used to annotate the resultant peptide (“peptides.txt”) files by organism and gene symbol. For each of the mouse and human tissue perspectives, data were analyzed two ways: 1) one in which peptides were included assuming only the organism of interest was present, and 2) one in which both organisms were assumed present for data normalization but separated into species-specific data sets. For method 1, ProteoClade filtered peptides into a data set in which the organism of interest only had to be a plausible assignment for each peptide and the other organism was excluded, simulating a targeted proteomics search in which only a single organism’s proteome was used as a reference. ProteoClade assigned peptide sequences and summed the MS2 intensities to gene symbols and then data were normalized by taking the relative intensities of each TMT channel compared to the internal reference pool for each TMT-plex used in the experiment, log2-transforming the data, centering the data by each channel’s median, and dividing the relative intensities of each channel by the channel’s standard deviation. For method 2, ProteoClade assigned peptides to organisms only if the peptide was unique to that organism in the context of the combined human and mouse proteomes. Downstream processing was similar to method 1, but the human and mouse genes quantified using species-unique peptides were separated to yield two distinct data sets after gene assignment and data normalization. Mouse and human data processing approaches were compared using the absolute difference between methods 1 and 2, and plotted using matplotlib 3.0.2.

To obtain a theoretical perspective of tryptic peptide similarity, the human and mouse Swiss-Prot (release 2018_01) reference databases were digested using a modified version of ProteoClade’s digest function. This yielded raw (i.e., non-compressed) peptide sequences which could be compared across homologs using their respective gene symbols.

### Differential gene expression of PDX stroma

The species-specific murine data set from Mundt et al.^2^ was selected for differential gene expression analysis. For each PDX WHIM sample, TMT channels corresponding to 2 hour vehicle, 50 hour vehicle, and “washout” were used as replicates. Genes were compared across WHIM samples using one-way analysis of variance (ANOVA), and the FDR was calculated and set to a limit of 0.05 using the Benjamini-Hochberg method (RStudio 1.0.153). Data were graphed using ggplot.

### Pathway analysis of PDX stroma

Differentially-regulated stromal gene lists from both PDX studies^2,3^ as determined by FDR-corrected ANOVA were combined for pathway analysis. Mouse genes were further filtered by removing plasma^4^ and abundant erythrocyte proteins identified by proteomics^5^ prior to downstream analysis. The web-based gene set analysis toolkit WebGestalt^6^ was used for overrepresentation analysis of the stromal proteins differentially regulated by PDX tumors in either dataset (2,2293 genes) versus those that were not (671 genes) using the default parameters, *Homo sapiens* as the organism, redundant datasets, and the affinity propagation option for redundancy reduction.

### Genus-level peptidome diversity

Taxonomic representation and sequence diversity at the tryptic peptide level was calculated using a modified version of ProteoClade’s PCDB module. A tryptic digest of the complete Swiss-Prot and TrEMBL sequences (release 2018_11) using ProteoClade’s default parameters was performed, but rather than assign peptides to organisms, non-compressed peptides were assigned directly to genera. Each peptide in the resultant database was checked for genus specificity, and the number of unique and shared peptides for each genus was tallied using custom scripts. Data for the number of unique peptides and fraction shared between genera were plotted using matplotlib.

### Oral microbiome *de novo* search

Raw spectra from the Grassl et al. study^7^ were retrieved from ProteomeXchange under ID PXD003028. A *de novo* search was performed using PEAKS Studio X (10) with the following parameters: the parent mass error tolerance was set to 10 ppm, the fragment mass error tolerance was set to 0.05 Da, the enzyme was set to trypsin, and MS2 fragmentation was set to higher energy collision-induced dissociation (HCD). Cysteine carbamidomethylation was set as a fixed modification and methionine oxidation was set as a variable modification. ProteoClade was used to make a PCDB of all 140.2 million Swiss-Prot and TrEMBL (release 2018_11) sequences for annotation using its default parameters as described in the implementation section.

### ProteoClade *de novo* annotation and genus identification

The searched results file (‘all de novo candidates.csv’) was filtered to only include candidate PSMs with a minimum average local confidence (ALC) score of 50. ProteoClade was used to annotate these PSM candidates using the “Database-Constrained” method, and assigned peptides to the species, genus, and superkingdom ranks. For the Database-Constrained method, candidates for each MS/MS spectra were ordered by descending confidence score, and the sequences were serially checked against the UniProt PCDB until a match, if any, was found. Peptides that did not belong to either to the bacteria superkingdom or human species were removed prior to further processing. Peptides with post-translational modifications were combined with their unmodified sequences for spectral counting, and peptides were only kept in the data set if they were unique to a single genus and had a minimum of two spectral counts across all samples. For FDR control, a PCDB with reversed protein sequences was created with ProteoClade, and the annotation steps were repeated against this database.

Genus-unique assignments from the *de novo* search that were compared to Grassl et al.’s original targeted database search if the genera were found in both data sets. Additionally, a comparison of ProteoClade’s top-scoring PSM annotation option to its Database-Constrained option was made. Genus-specific assignments were ranked by taking the difference between their forward and reversed PCDB annotations. Plots were made using Plotly.

---

**Figure S1:**
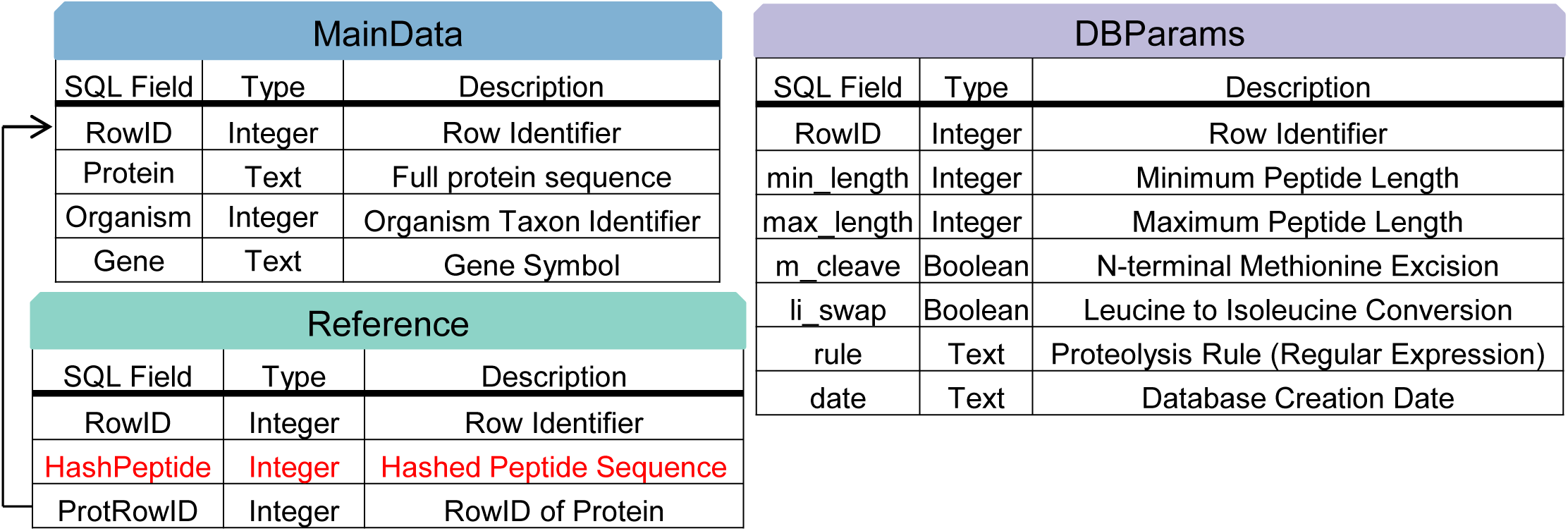
Database schema for the ProteoClade Database (PCDB). The database stores information across three tables: “MainData” stores complete protein sequences, organisms, and genes; “Reference” contains all peptide information and a key back to the protein table (black arrow); “DBParams” stores all database parameters at the time of database creation. The indexed column, “HashPeptide,” is indicated in red.

**Figure S2:**
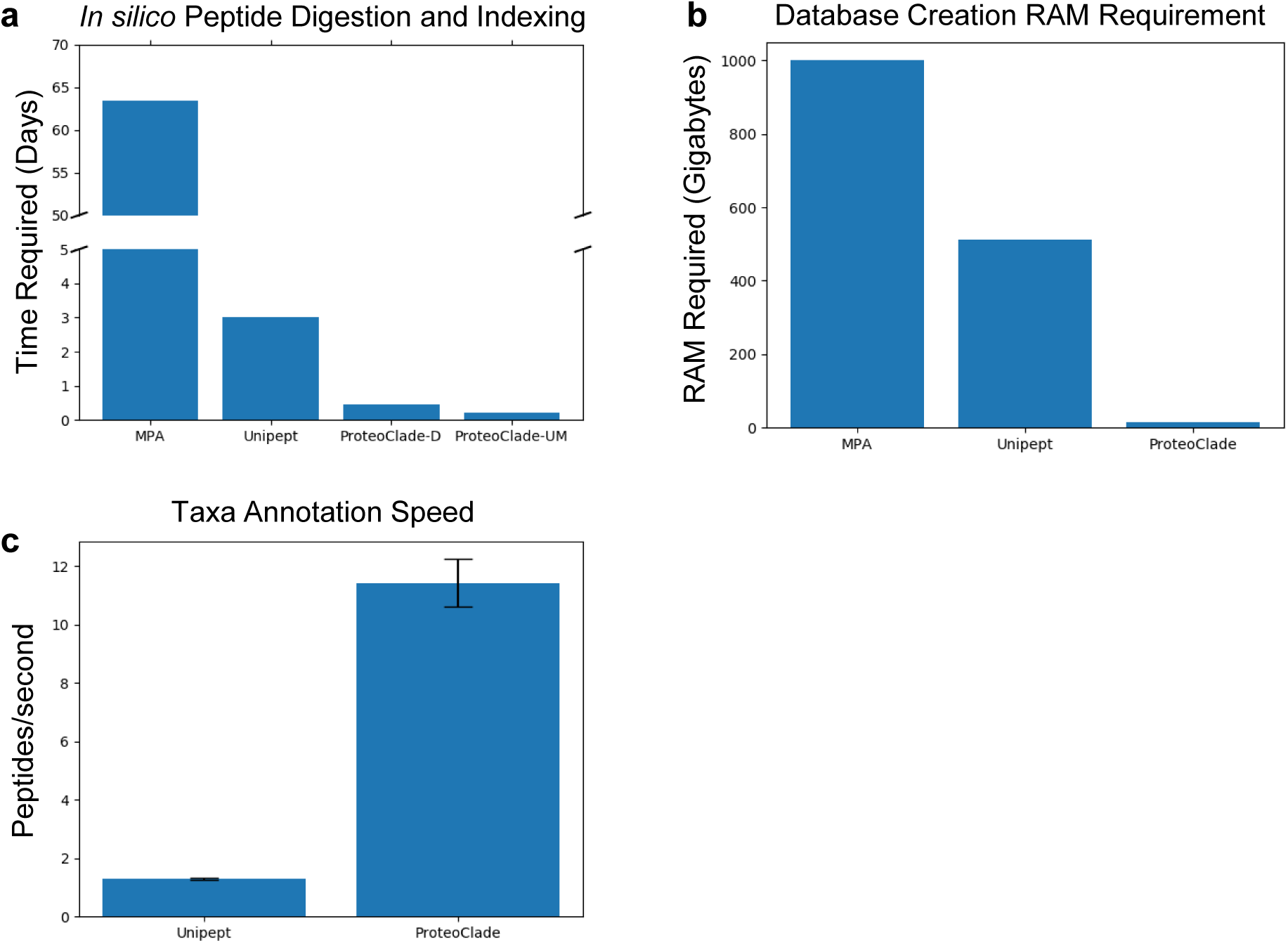
Technical comparisons between metaproteomic analysis tools. **a)** *In silico* digestion of peptides and indexing for ProteoClade’s default settings (ProteoClade-D) is faster than previous tools when using the entire UniProt repository. A database using the same parameters as Unipept (ProteoClade-UM) was faster still, due to the absence of missed cleaved peptides. **b)** Database RAM requirements for ProteoClade enable users to generate large databases without using high performance computers. **c)** Annotating all taxa for experimental results is 8.8x faster for ProteoClade than Unipept.

